# Genome-wide fluctuating selection under demographic cycling in Icelandic rock ptarmigan

**DOI:** 10.64898/2026.10.28.762602

**Authors:** Alexandre Rêgo, Theodore E. Squires, Wanyi Wei, Patrik Rödin-Mörch, Jacob Höglund, Kristinn P. Magnússon

## Abstract

Understanding how natural populations adapt to continuously shifting environments is central to evolutionary biology. Multi-annual population cycles impose temporally varying selective pressures, yet disentangling the resulting genomic signals from those of sustained directional selection and genetic drift has been challenging due to a lack of high-resolution temporal data. Here we investigate the genomic consequences of demographic cycling in a population of rock ptarmigan (Lagopus muta) from northeastern Iceland. We analyzed whole-genome sequences from 91 individuals sampled over an 11-year period, encompassing two population peaks (2010 and 2018) and two troughs (2007 and 2013). Testing allele frequency change at each SNP against a null model of drift and sampling, we identified candidate loci and grouped them by correlation of their temporal trajectories. Cluster trajectories fluctuated through time, and two were closely correlated with the regional census, one significantly tracking the demographic cycle, while a smaller set changed monotonically in a manner consistent with sustained directional selection. Each cluster contained loci distributed across many chromosomes, indicative of genome-wide responses to selection. The most strongly fluctuating loci overlapped candidate genes of possible functional relevance, most notably CTNNA2 (Catenin Alpha-2), which is associated with startle response and anxiety-like behaviors and may relate to predator avoidance. Our findings indicate that fluctuating selection comprises much of the temporal genomic response in this natural population, and suggest that multi-annual demographic cycles can drive rapid and widespread allele frequency change, highlighting the value of temporal genomics for clarifying the interplay between demography and natural selection in wild populations.

## Introduction

The magnitude and direction of natural selection in nature are very rarely constant, even over short ecological timescales. In response to temporally variable environments, populations may track the environment through changes in allele frequency at polymorphic sites across the genome (i.e., adaptive tracking). The extent to which alleles track shifting conditions has been difficult to establish, however, because low-resolution temporal sampling is unable to resolve allele frequency trajectories, and selection on standing variation is often modest and can resemble genetic drift (Messer et al. 2016; Buffalo and Coop 2020).

When environmental shifts are regular and cyclical, as in seasonality or when frequency and density dependence act at the demographic level, the strength and direction of selection can vary in step with the environment, so that alleles favored under one set of conditions become disfavored under another. Such fluctuating, temporally varying selection can produce coordinated changes in allele frequency that rise and fall with the environment (Bell 2010; Lande 2007). Much of the direct evidence for this comes from Drosophila, where seasonal cycles drive strong parallel allele frequency changes across many loci and replicate populations over only a few generations (Rudman et al. 2022; Bitter et al. 2024; Machado et al. 2021); at seasonally oscillating loci, allele frequencies shifted on average by 20% in as few as 2 generations (Bergland et al. 2014).

The interactions between demography and selection nonetheless remain poorly resolved (but see (Nespolo 2007; Gulisija and Kim 2015; Jain and Kaushik 2022). A central and largely open question is how much of the genomic response to a changing environment reflects fluctuating selection that reverses as conditions cycle, as opposed to sustained directional selection (Soni and Jensen 2024). The two processes leave different temporal signatures, with fluctuating selection producing frequencies that rise and fall alongside the environmental driver and directional selection producing monotonic change, yet both can act at once on different loci within a single population. Separating their relative contributions requires sampling allele frequencies repeatedly through an environmental cycle, since a single interval cannot distinguish a reversal from one step in a longer trajectory. Quantifying the relative contribution of each would clarify how much contemporary genomic change represents transient tracking of a shifting optimum rather than lasting adaptation.

In Iceland, the rock ptarmigan (Lagopus muta) provides an excellent opportunity to investigate selection over time with influences from complex demography. Icelandic rock ptarmigan populations are well-studied with extensive historic sampling data available (Garðarsson 1971, 1988), and the present study population shows well-documented multi-annual demographic cycling (Nielsen 1996; Sturludóttir et al. 2018; Johnson and Nielsen 2024). These cycles are large in amplitude (as much as twenty-fold between troughs and peaks) and recur with a variable period on the order of four to ten years. This cyclicity has been linked to density-dependent predation by the gyrfalcon (Nielsen and Pétursson 1995) and endoparasite load (Skírnisson et al. 2012); both operating when high prey/host density elevates predation or parasite transmission, in turn depressing survival and fecundity and feeding back on the decline. These cycles have also been associated with concomitant shifts in phenotypic traits (e.g., fat, gut length) (Guðmundsson 2015; Stenkewitz et al. 2016). These regularly shifting ecological conditions create a setting in which alleles at many loci may alternate between beneficial and deleterious depending on whether the population is in a peak or trough (Botero et al. 2015; Himes Boor et al. 2018; Rudman et al. 2022), making the ptarmigan a tractable wild system in which to look for adaptive tracking paired with cycling demography.

Using whole genome re-sequencing data of a single population of Icelandic rock ptarmigan at four different time points across an 11-year period, spanning two demographic troughs and two peaks, we ask whether allele frequencies track the demographic cycle and characterize the genomic architecture of any such tracking. We test whether sets of loci across the genome share correlated temporal trajectories, whether those trajectories align with the independently monitored regional census, whether any co-fluctuation is genome-wide or restricted to a limited number of loci, and the relative contribution of fluctuating and directional selection in this system. We further identify candidate genes underlying the most strongly fluctuating loci and consider their possible functional relevance.

## Methods

### Sampling of Ptarmigan

Rock ptarmigan individuals were collected each autumn as part of the comprehensive “Rock ptarmigan health and population change” project to investigate the species’ health status in Iceland between 2006 and 2018 (Stenkewitz et al. 2016). We utilized a subset of 91 available young-of-the-year birds from these studies collected from a 150,000-hectare area of northeastern Iceland located between Lake Mývatn and Húsavik to ensure capturing of local demographics (Fig. 1). We utilized spring census data of the Rock Ptarmigan collected via transect-based surveys (Johnson and Nielsen 2024; Ferrarini and Nielsen 2025). The northeastern region has recently seen spring ptarmigan census sizes fluctuating between 10,000 and 55,000 birds and shows the strongest cyclicity in the country (Johnson and Nielsen 2024; see Fig. 2). We sampled individuals from years representing both population troughs (2007, census = 22600, n = 26; and 2013, census = 23400, n= 24) and peaks (2010, census = 51400, n = 20; and 2018, census = 30800, n= 21). Samples included both females and males at an approximately 1:1 ratio each year (F_07_ = 11, M_07_ = 15; F_10_ = 10, M_10_ = 10; F_13_ = 13, M_13_ = 11; F_18_ = 9, M_18_ = 12; F_all_ = 43, M_all_ = 48). We note that two of the four years are offset by one year from the demographic extremes (2013 one year after the trough and 2018 one year before the following peak) due to choosing sampling years with the strongest trough and peak signals across the Icelandic-wide population rather than the local extremes (Johnson and Nielsen 2024). Further, the census tracks breeding adults each spring, whereas the genotyped birds are their young-of-the-year offspring collected that autumn. Thus, the allele frequencies of the genotyped cohort reflect selection under the reported census size.

**Figure 1.**
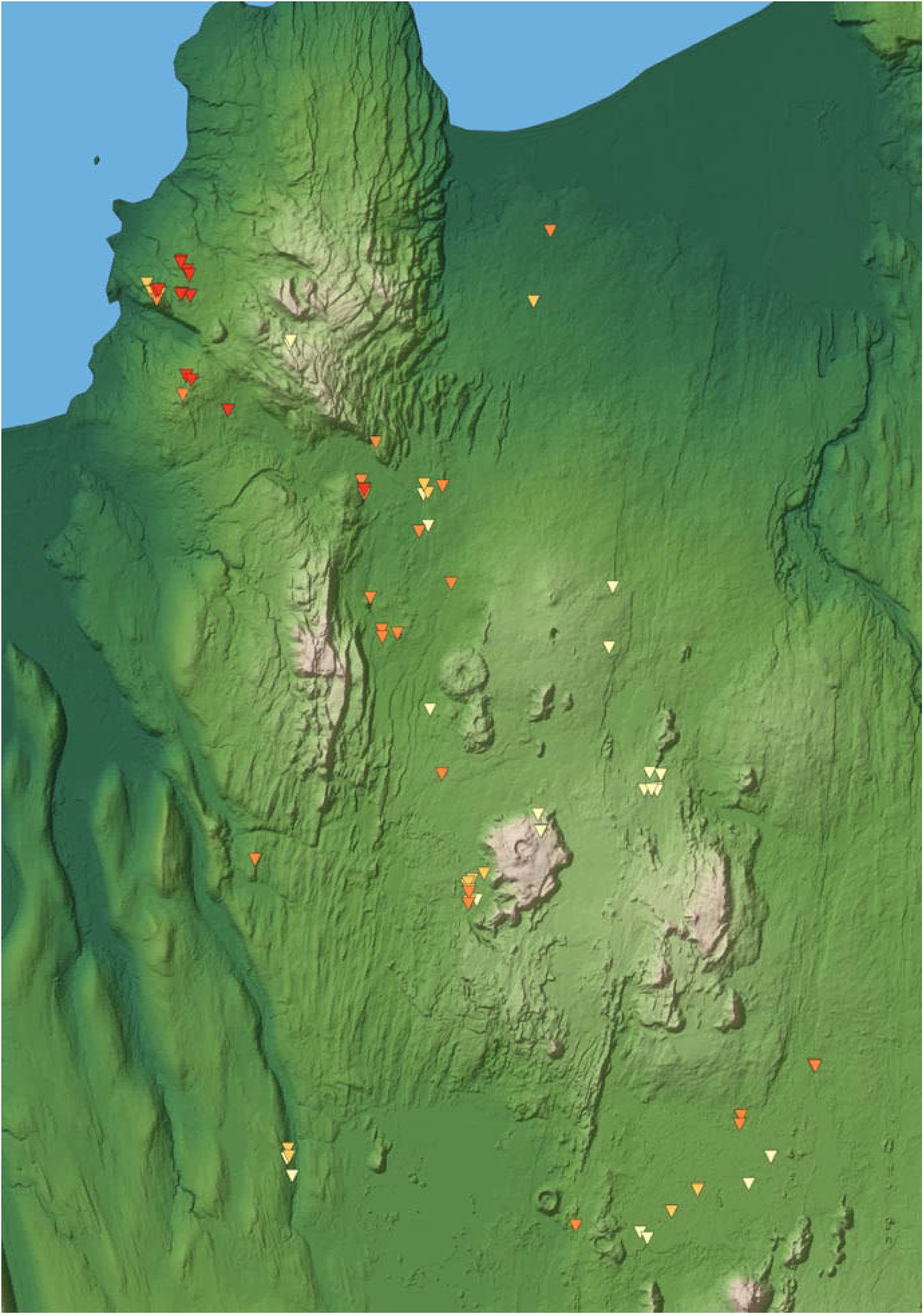
Relief map showing the years and locations of Rock Ptarmigan used in the present study from around northeastern Iceland approximately between Husavík and Reykjahlíð. The number of sampled individuals from the various years are 2007, n = 26 (light yellow); 2010, n = 20 (orange); 2013, n = 24 (dark orange); and 2018, n = 21 (red).

**Figure 2.**
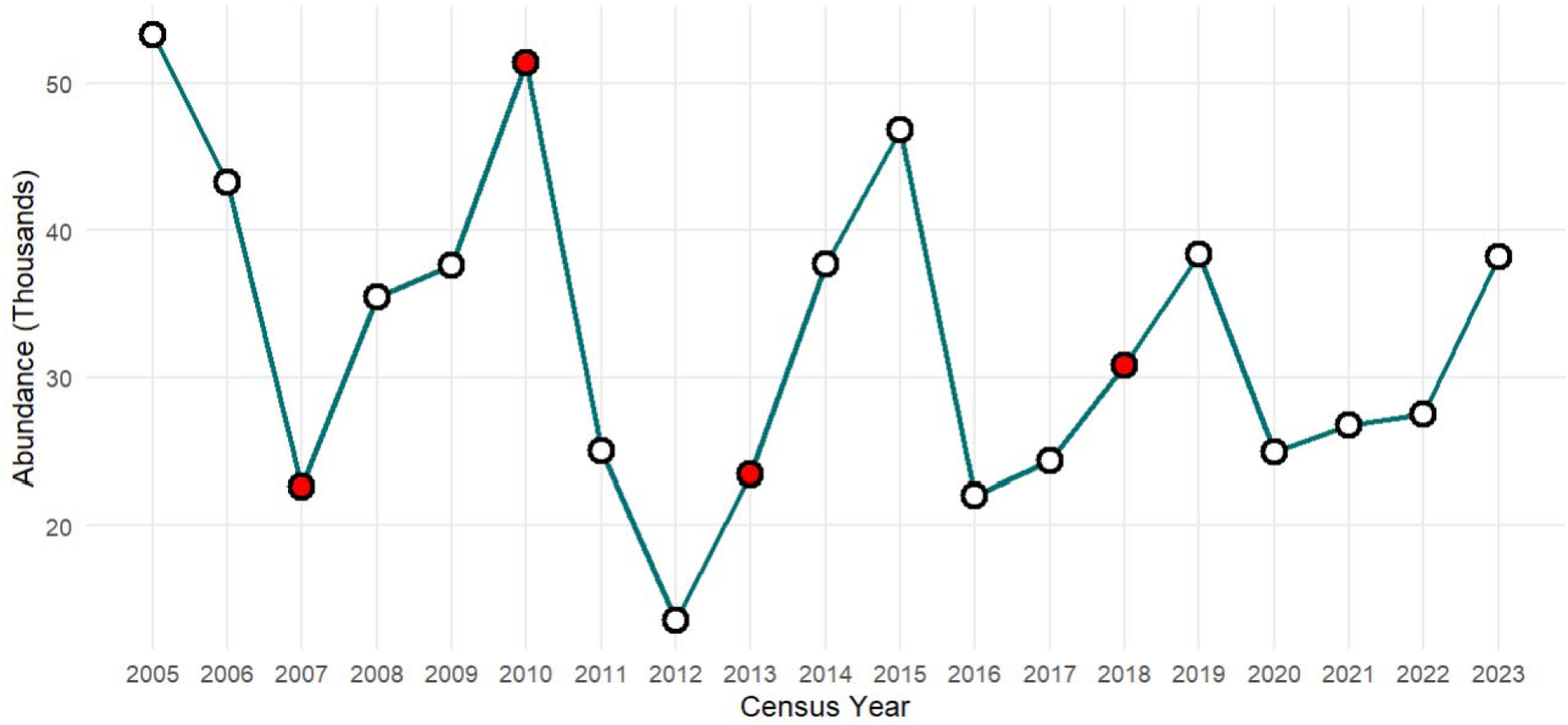
Overview of spring population census data from rock ptarmigan counted in the Northeastern Region (Norðurland eystra) of Iceland between 2005 and 2023 as collected by the Natural Science Institute of Iceland (methods detailed in Ferrarini and Nielsen 2025). Our sampling years are highlighted with bold red points. This figure was developed from data previously reported by Johnson and Nielsen (2024) and made available in their dryad repository (https://doi.org/10.5061/dryad.c59zw3rg3).

DNA was extracted from muscle tissue was carried out in a dedicated sterile genetics lab using established protocols from a Monarch Genomic DNA Purification Kit (New England BioLabs, USA) and quantified for concentration and purity on a NanoDrop 2000 spectrophotometer and a Qubit 3.0 fluorometer (Invitrogen). DNA Nanoball library preparation and DNBSEQ PE150 paired-end whole-genome sequencing at approximately 20x coverage were performed by the Beijing Genomics Institute Group (Hong Kong).

### Preparation of Genomic Data

Read quality was assessed with FastQC v.0.11.9 (Andrews 2010), and adapters and low quality sites were removed with Trimmomatic v.0.36 (Beier et al. 2026). Trimmed reads were aligned to the rock ptarmigan reference genome (GenBank accession: GCA_023343835.1; (Squires et al. 2023) using BWA-mem v.0.7.17 (Li and Durbin 2010), sorted and duplicate-marked with Picard v.2.20.4 (Broad Institute, 2019), and assessed for mapping quality and coverage with Qualimap v.2.2.1 (Okonechnikov et al. 2016). Variants were called using GATK v.4.1.1.0 (McKenna et al. 2010) following best practices (DePristo et al. 2011; Van der Auwera et al. 2013), running Haplotypecaller in GVCF mode per individual and joint-genotyped with GenotypeGVCFs.

We retained biallelic SNPs and applied GATK best-practice hard filters, removing sites with QUAL < 30, MQ < 40, QD < 2, FS > 60, SOR > 4, MQRankSum < −12.5, or ReadPosRankSum < −8. We then filtered by read depth (10 < DP < 41), allowed up to 10% missing data, removed SNPs within 5 bp of an indel, and required a minimum minor allele frequency of 0.05 across the pooled samples, resulting in a final set of 7,174,144 SNPs.

Relatedness estimated in VCFtools (Danecek et al. 2011), using both the A_jk_ statistic and KING-based coefficient (Φ; Manichaikul et al. 2010), revealed no close relatives. Only one pair of individuals showed any potential relationship (LM-13-122 and LM-18-052B; A_jk_= 0.114, Φ= 0.099), although this was at a level below first-cousins.

PrincipalTcomponent analysis of individual genotypes at all autosomal SNPs (chromosome 16 excluded, as its large inversion polymorphism otherwise dominates the leading PCs) showed no separation of individuals by sampling year, indicating no detectable temporal population structure (Fig. S1). A Mantel test of pairwise genetic against geographic distance (Pearson r = 0.057; 9,999 permutations, oneTsided p = 0.106) found no association over the ∼60 km sampling area, consistent with a single panmictic population and supporting the pooling of individuals across sites (Fig. S2).

### Calculating Genetic Diversity (π), differentiation (F_ST_), and allele frequencies

Autosomal nucleotide diversity (π) and pairwise differentiation (F_ST_) were computed utilizing pixy v.2.0.0.beta14 (Korunes and Samuk 2021) over 10kb sliding windows. To identify genomic regions differentiating with the population census cycle, we contrasted pooled trough years (2007 and 2013; n = 50) against pooled peak years (2010 and 2018; n = 41), estimating Z-standardized F_st_ (ZF_st_) per window after excluding sparse windows (<10 SNPs). Windows exceeding the 99th ZF_ST_ percentile were classified as outliers (913 windows). Gene annotations were obtained directly from NCBI (Assembly ID: GCF_023343835.1), considering autosomes only, resulting in 18,872 annotated genes.

### Identifying loci under selection

To identify loci under fluctuating selection, we first identified candidate loci consisting of putatively selected SNPs. For each SNP we tested the observed change in allele frequency against a null model of genetic drift and finite sampling, evaluated between each consecutive sampling year (2007–2010, 2010–2013, 2013–2018) and the full 2007–2018 span. We first estimated the contemporary variance effective population size (Ne) between each time interval from patterns of allele frequency change from the total set of 7,174,144 SNPs using the software `varne` (Gompert and Messina 2016; Rêgo et al. 2019). These estimates were subsequently used to parameterize the null model of expected evolutionary change due to genetic drift. These interval estimates reflect the effective number of breeders across each interval rather than the census size at either endpoint, so they need not track the census at the sampled years. Due to our modest sample size at each time point (2N = 40–52), we also account for the effects of sampling error in our null models.

The null model combines three sources of variance (Fig. S3): uncertainty in the true initial frequency given the observed count _0_ of _0_ sampled gene copies, 0∼(_0_ + 0.5, _0_ - _0_ + 0.5); genetic drift over t generations, which under a beta approximation to the Wright–Fisher process (Ewens 2004; Gompert and Messina 2016) p_0_(1 - p)F adds variance, where 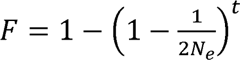; and binomial sampling of _1_ gene copies at the final timepoint. These combine in closed form into a beta-binomial distribution for the final allele count, _1_∼ - (_1_,) with 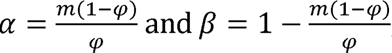, where 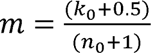 is the expected initial frequency and 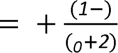 is the overdispersion of the beta-binomial, combining drift with the uncertainty in the initial allele frequency. From this distribution we computed a two-sided p-value as the total probability of outcomes no more likely than the one observed, and retained a SNP as a candidate if it reached p<0.0001 in any of the time-interval comparisons, yielding 2,925 candidate SNPs. It should be noted that SNPs identified in this manner are not necessarily causal loci, but are likely in linkage disequilibrium with causal loci, and are representative of the patterns of evolutionary change in those loci. We validated this closedTform betaTbinomial test against a 200,000Tdraw Monte Carlo simulation of the same threeTstage model (pTvalues fall on the 1:1 line) and confirmed that its falseTpositive rate tracks the significance threshold across a range of true starting frequencies (Fig. S4).

To describe the overall allele frequency dynamics in putatively selected loci, we constructed clusters of SNPs based on correlations between allele frequency trajectories through time (Otte and Schlötterer 2021), Ament et al. 2022). Allele frequencies of the 2,925 candidate SNPs were normalized with an arcsine-square-root transformation followed by centering and scaling. We calculated Pearson correlations between the entire subset of SNPs, converted these to distances (1 - |r|), and performed hierarchical clustering using average linkage with a maximum distance threshold of 0.2. A total of 4 clusters were observed with at least 10 SNPs in the cluster. To assess whether these clusters tracked changes in population size, we compared each cluster’s mean allele-frequency trajectory to reported regional census counts at the four sampling years. For each cluster we computed the Pearson correlation (|r|) between its mean trajectory and the census series across the four timepoints, and assessed significance with an exact permutation test.

## Results

### Diversity and Differentiation Among Sampling Years

Within-year nucleotide diversity was stable across the study (π ≈ 3.18 x 10^−3^), reflecting long-term effective population size rather than contemporary census shifts, and genome-wide differentiation between phases was negligible (mean F_ST_ = 3.6×10⁻⁵), consistent with limited genetic drift over only a few generations. Given this low differentiation, ZF_ST_ windows here represent a relative ranking of the most differentiated windows rather than a formal test of selection; high ZF_ST_ windows were treated as candidates and cross-referenced against the candidate clusters of the allele frequency trajectory analysis (see below). The top 1% of windows (913 windows), overlapped 537 annotated genes. Elevated ZF_ST_ windows were uncorrelated with gene density (point-biserial r = −0.007) such that outlier windows are not reflective of gene-dense regions of the genome. The 537 genes overlapping the top-1% windows represent 2.8% of the 18,872 annotated autosomal genes.

Among the top 10 most strongly differentiated windows (Fig. 3), the top-ranked window genome-wide overlapped with CTNNA2 (ZF_ST_ = 21.6), with a second, adjacent CTNNA2 window also among the top ten. Further outlier windows overlapped RPA3/UMAD1 (two adjacent windows on Chromosome 7), PPP1R13B, RPS3A/SH3D19, ATP10D, and the long non-coding RNA LOC125692110, while two intergenic outliers lay nearest to MARCKS and TRAF3; five of the ten strongest windows fell on chromosome 4. GO enrichment of the 537 ZF_ST_ candidate genes recovered nine significant synaptic cellular-component terms (FDR<0.05), including glutamatergic synapse, postsynaptic density, and synaptic membrane, consistent with the lead candidate CTNNA2 (Fig. S5).

**Figure 3.**
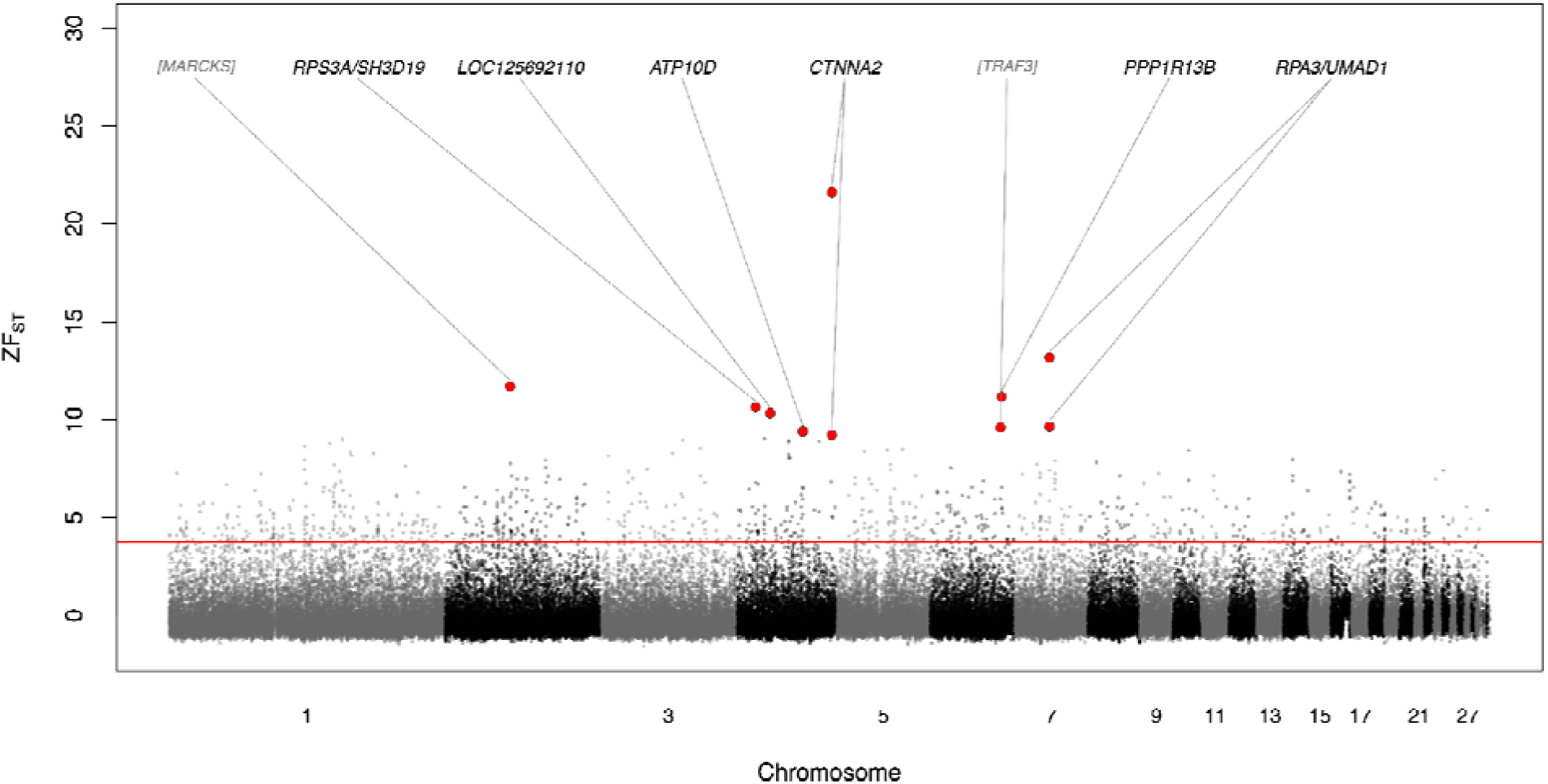
Differentiation (ZF_ST_) among SNPs in 10 kB non-overlapping windows between pooled peak (2010 and 2018) and trough years (2007 and 2013). Protein coding genes in the ten top outlier windows are indicated, with nearest genes in brackets for windows with no genes present. The red line indicates the top 1% quantile.

### Characterizing fluctuating selection

The effective population sizes associated with each time interval were large, with moderate variation (Ne_07-10_ = 2,878 [95%CI: 2539.06, 3400.25]; Ne_10-13_ = 897 [858.623, 940.793]; Ne_13-18_ = 5633 [4803.08, 6921.54];Ne_07-18_ = 1,261 [1236.61, 1288.37]). This variation is unsurprising, as the census population size itself fluctuates rapidly. Despite this variation, Ne is sufficiently large between each time interval that we expect genetic drift to contribute relatively little to changes in allele frequency (Fig. S6).

To summarize allele frequency dynamics across putatively selected loci, we clustered the 2,925 candidate SNPs (p < 0.0001 in any of the four time intervals) based on correlations in their temporal trajectories (Fig. 4). Four clusters were recovered (646–925 SNPs), which together comprised all candidate SNPs. Each cluster contained SNPs from at least 27 different chromosomes (Fig. S7), indicating that correlated allele frequency change is distributed across the genome rather than single linked regions. Within every cluster, linkage disequilibrium decayed to the genomeTwide background (r² ≈ 0.05) by ∼100 kb in all four sampling years, confirming that cluster members more than ∼100 kb apart are effectively unlinked and that the shared trajectories are not a product of physical linkage (Fig. S8). Clusters generally exhibited non-linear patterns of allele frequency change, with oscillating or stepwise trajectories. Two clusters in particular (Fig. 4A and 4B) exhibited allele frequency changes which were highly correlated with fluctuations in demography of the ptarmigan populations (cluster-mean |r| with census = 0.90 and 0.89; Fig. S9), with cluster A significantly tracking the census across the four sampled years (p = 0.042). One cluster in particular (Fig. 4C) exhibited broadly monotonic increases in allele frequency on average (38.8% of SNPs strictly increasing every year), consistent with sustained directional selection over the course of sampling.

**Figure 4.**
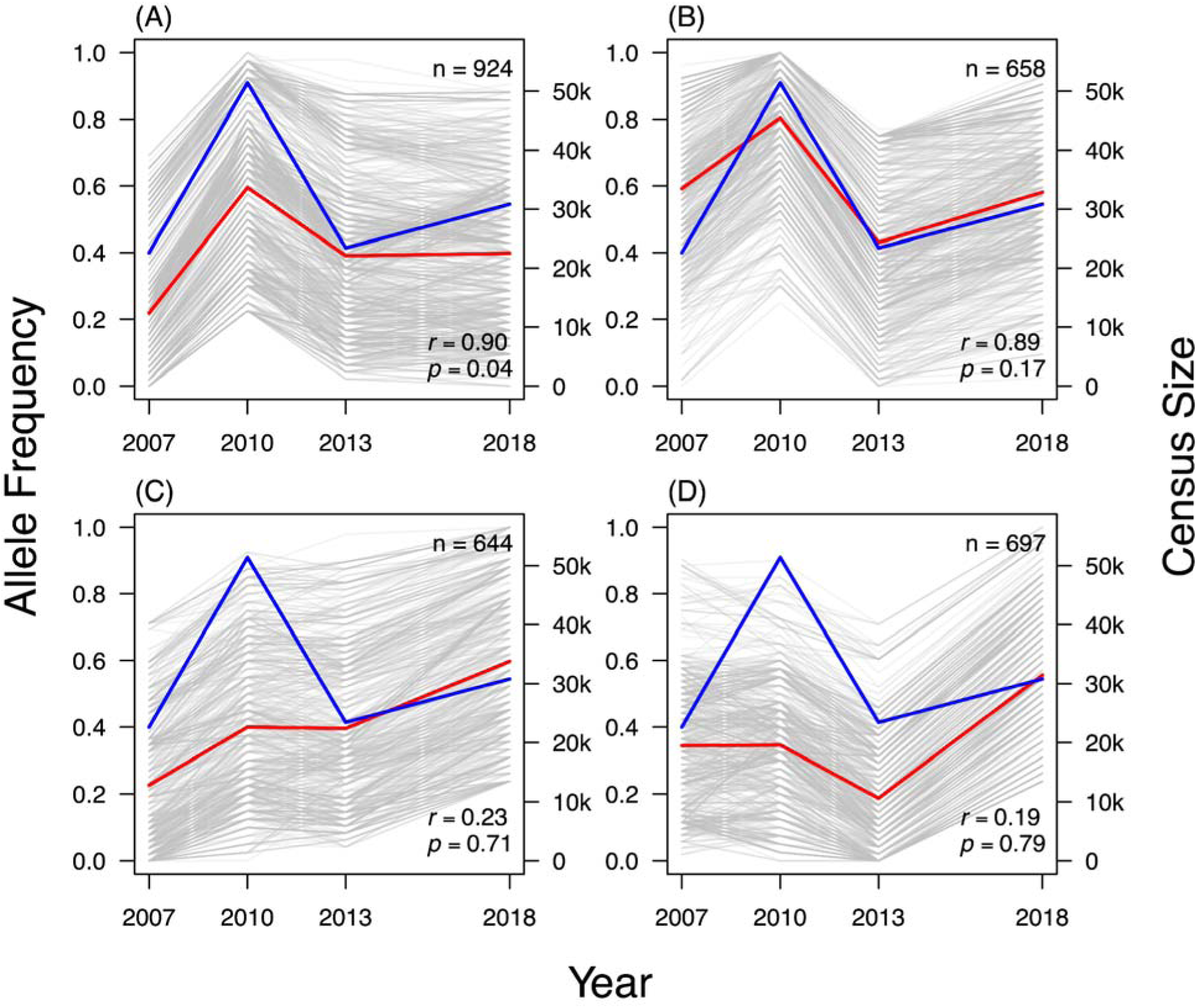
Clusters of allele frequency changes in putatively selected loci. Individual frequency trajectories are shown in grey, the mean allele frequency of the cluster is in red, and the census trend is shown in blue. The number of SNPs contained within each cluster is shown (of the 2,925 candidate SNPs which showed p < 0.0001 in any time interval), as well as the correlation between the mean allele frequency (red) and census size (blue). Note that nearly all clusters exhibit patterns of fluctuating selection of some form (i.e., non-linear patterns of allele frequency change). For each SNP the allele plotted was chosen to align with the cluster mean, and clusters were oriented with the census; absolute allele identity is arbitrary. Regional census data as previously reported in Figure 2.

Enrichment differed markedly between the census-tracking and non-tracking clusters. Surprisingly, the two clusters whose trajectories tracked census size (clusters A and B) carried little functional signal. Cluster A was enriched only for broad cellular-component terms (plasma membrane, cell periphery; FDR < 0.05), and cluster B yielded no significant terms. In contrast, the two clusters that did not track census were enriched for cell-adhesion and signaling functions. Cluster C, which showed sustained directional change, was most strongly enriched for immunoglobulin-superfamily cell-adhesion-molecule (IgSF CAM) signaling (KEGG; FDR = 9 × 10⁻⁵), along with C-type lectin receptor, calcium, VEGF and MAPK signaling pathways and voltage-gated ion-channel activity (Fig. 5). Cluster D was enriched for phosphoinositide metabolism and signaling (inositol phosphate metabolism, phosphatidylinositol dephosphorylation, phosphatidylinositol signaling) and, like cluster C, for IgSF CAM signaling.

**Figure 5.**
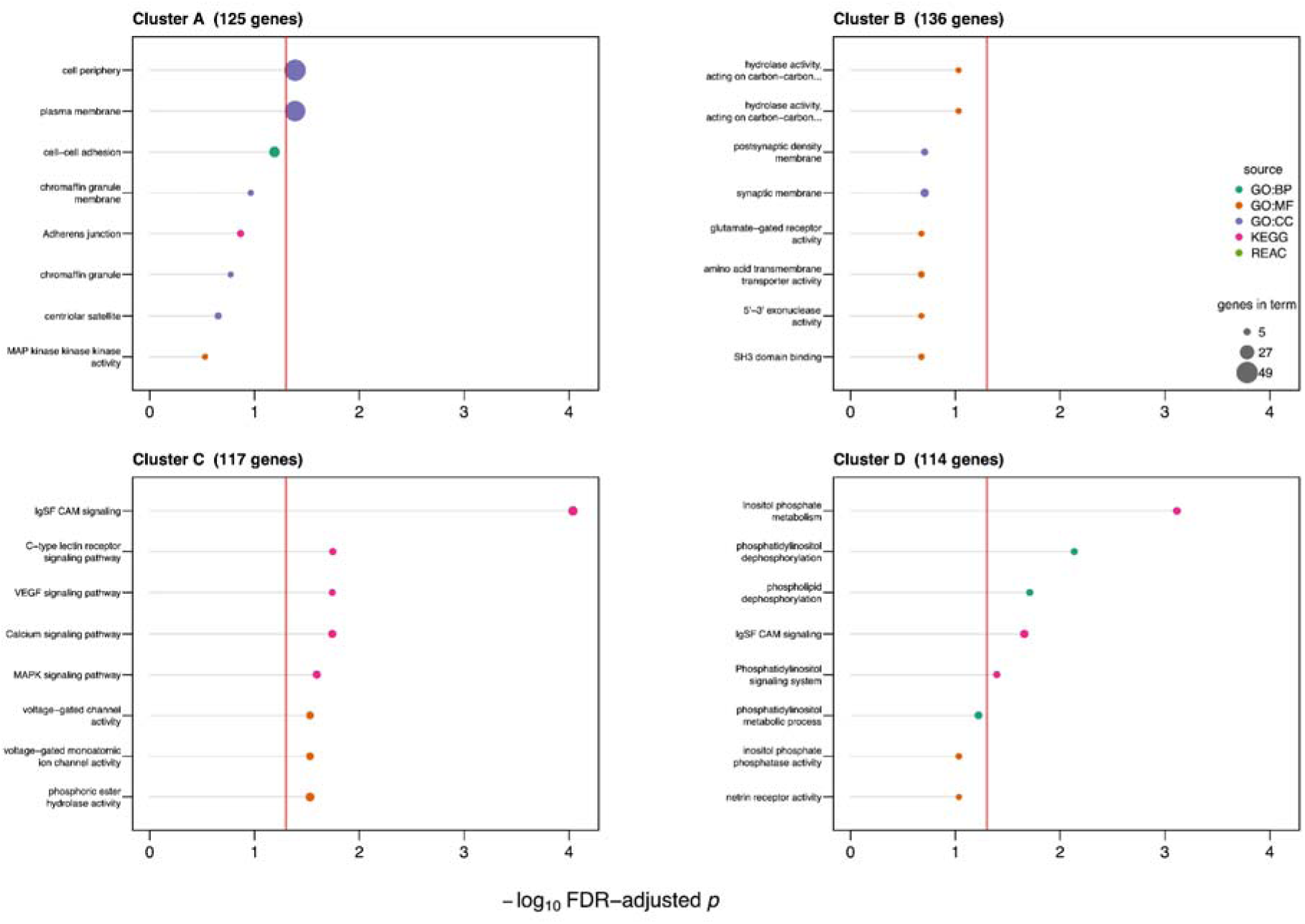
Functional enrichment of the trajectory clusters. Genes overlapped by the SNPs in each trajectory cluster (A–D; number of protein-coding genes per cluster given in the panel headers) were tested for Gene Ontology (biological process, molecular function, cellular component) and pathway (KEGG, Reactome) enrichment using g:Profiler, with Gallus gallus orthologues and all named protein-coding genes genome-wide as the background, and Benjamini–Hochberg FDR correction. Each panel shows the eight most enriched terms for that cluster, ordered by significance; the vertical red line marks FDR = 0.05. Point color indicates the annotation source and point size the number of cluster genes contributing to the term (common scale across panels).

Despite few functional signals being associated with cluster A, the cluster that best tracked census size, several of the highest-ZF_ST_ genes (CTNNA2, RPA3 and UMAD1) also fell within it. While not completely independent tests, these approaches are nonetheless complementary; pooling phase-matched years gives ZF_ST_ greater power to detect and rank the magnitude of peak–trough differentiation, but discards the temporal order and the direction of change, whereas the clustering retains the full four-timepoint trajectory and correlates it against independent census data which both the ZFST scan and candidate-detection step are agnostic to.

## Discussion

We find that fluctuating selection comprises much of the temporal genomic response in this cycling population of rock ptarmigan. Testing allele frequency change at each SNP against a null model of drift and sampling recovered 2,925 candidate loci whose trajectories, when clustered by their temporal correlation, formed four groups each distributed genome-wide. Two of the four clusters were closely correlated with the regional census, and one tracked it significantly across the four sampled years. Several lines of evidence argue against drift as the source of these patterns. Effective population sizes between intervals were large and genome-wide differentiation between phases was negligible. Taken together, the coordinated, genome-wide census-correlated changes are most consistent with selection that varies in sync with the demographic cycle. It is important to note that this inference does not require the sampled years to fall on the exact demographic extremes; each cluster is correlated against the census count record at each sampled year which still describe an extreme fluctuating non-monotonic density series that a set of loci can correlate with only by rising and falling in step with it.

The architecture we observe, with selection spread across many co-varying loci is similar to seasonal adaptive tracking documented in Drosophila (Rudman et al. 2022; Bitter et al. 2024; Machado et al. 2021; Bergland et al. 2014), and genome-wide fluctuating selection recently reported in Daphnia (Lynch et al. 2024). Fluctuating selection has also been demonstrated in wild stickleback populations (Gasterosteus aculeatus), though with transient cycles of variable length (Phillips et al. 2023; Strickland et al. 2024). Notably, the clearest signals of fluctuating selection so far come from intra-annual cycles in invertebrates that pass through many generations per environmental period, whereas the ptarmigan tracks a multi-annual cycle across only a few generations per period. The degree to which a population tracks its environment genetically, rather than through phenotypic plasticity, likely depends on its generation time relative to the period of environmental change (Chevin et al. 2010; Botero et al. 2015).

The cyclicity that provides the environmental driver is itself well documented. Icelandic rock ptarmigan populations fluctuate as much as twentyfold between troughs and peaks, with a variable period on the order of four to ten years (Garðarsson 1988, Nielsen 1996, Sturludóttir et al. 2018, Johnson and Nielsen 2024). The cycle has long been attributed to density-dependent predation by the gyrfalcon, the principal predator of the ptarmigan (Nielsen and Pétursson 1995), with parasite load and transmittance increasing with density proposed as an additional factor (Stenkewitz et al. 2016; Johnson and Nielsen 2024). As a game species, ptarmigan is hunted in Iceland for personal consumption, although the annual take has been roughly constant in recent decades and isn’t understood to exceed sustainable harvest or drive demographic cycles (Johnson and Nielsen 2024). Importantly, first year birds are often collected in the fall when hunters cannot easily distinguish them from adults. It is therefore probable that harvest is an additional selective agent on juvenile cohorts alongside predation and parasitism, and could warrant further investigation. Rock ptarmigan colonized Iceland following deglaciation roughly 10,000 years ago (Holder et al. 1999; Sahlman et al. 2009), so the population has been exposed to recurring, density-linked shifts in predation and condition over many thousands of generations. It is plausible that selection regimes fluctuating with density have shaped standing allelic variation over this time, whether through effects that reverse in sign as density changes or through frequency-dependent dynamics, though these mechanisms remain candidate explanations.

Among the candidate genes, CTNNA2 stands out as having been found in the single most differentiated ZFst window and also fell within the census-tracking cluster A. CTNNA2 encodes alphaN-catenin, which is expressed in the developing brain, and has been linked to the control of startle responses in mice (Park et al. 2002) and to anxiety-related phenotypes in humans (Eszlari et al. 2021). Flushing behavior, which is closely related to startle, decreases with population density in ptarmigan (Pelletier and Krebs 1998), and predation risk from the gyrfalcon changes as density cycles, so an allele influencing startle or vigilance could plausibly be favored differently between demographic peaks and troughs. More broadly, the census-tracking clusters lean toward neurological and behavioral functions at the level of individual genes, including glutamate receptor subunits and other synaptic components. This is consistent with the possibility that behavioral responses to changing density and predation form part of what tracks the cycle, though we treat this as a hypothesis rather than a demonstrated mechanism. Density-dependent shifts in social structure and behavior have been documented in the closely related red grouse (Piertney et al. 2008; Cattadori et al. 2005).

In contrast to the census-tracking clusters, the clusters that did not track the census were enriched for immune and signaling functions, including immunoglobulin-superfamily cell-adhesion-molecule and C-type lectin receptor signaling in the directional cluster and phosphoinositide signaling in the remaining cluster. The parasite Eimeria is common in Icelandic rock ptarmigan and has been proposed as a contributor to the cycle (Stenkewitz et al. 2016; Johnson and Nielsen 2024), so it is tempting to read an immune signal as parasite-driven selection. In our data, however, the enriched immune-related terms fall in the clusters that do not track density, and individual immune-annotated genes are scattered across clusters rather than concentrated among the strongly fluctuating, census-tracking loci. We therefore do not interpret specific gut-immunity genes as fluctuating with density, though parasite-mediated fluctuating selection may act at a different periodicity than sampled here.

While the dominant mode of selection among candidate loci is fluctuating selection, one cluster increased monotonically, with 38.8% of its SNPs rising at every timepoint, consistent with sustained directional selection. Sustained directional change over the sampling period could reflect environmental drivers operating on timescales longer than the cycle, including the rapid warming of the Arctic, where rising temperatures, reduced snow cover, and greater seasonality of rainfall are expected to alter selective pressures on ptarmigan (Post et al. 2009; Box et al. 2019; Höglund et al. 2019; IPCC 2023). At the same time, an apparently directional trajectory sampled over 11 years may be part of a fluctuation whose period exceeds our window. These explanations are not mutually exclusive.

The questions that remain are, in large part, questions of temporal resolution. Whether the loci we classify as directional are under sustained selection or are tracing slow fluctuations, and whether any of the fluctuating variation we observe is maintained over the long term rather than simply turned over, cannot be settled within a single 11-year window. A longer series could show whether alleles that rise and fall with density return to comparable frequencies across successive cycles, and such a signal of return, sustained over many cycles, would begin to connect the rapid fluctuating selection described here to the long-term maintenance of polymorphism that defines balancing selection, a link that temporal data of sufficient depth are uniquely placed to test. Whole-genome time series of naturally cyclic wild populations thus offer a direct route to questions about the interplay of demography and selection that cross-sectional data leave unresolved.

## Acknowledgements

We wish to thank surveyors and hunters with the Ptarmigan Population Health Project (Vöktun rjúpnastofnsins) and material handlers from the Natural Science Institute of Iceland. Specifically, Ólafur Karl Nielsen was invaluable for material provisioning along with providing much of the theoretical background for this work. We wish to thank Athanasios Toros for reviewing the integrity of our genomic data, and Uppsala University undergraduate Johanna Ehlton for assistance with figure development in GIS. This study was financially supported by the Swedish Research Council (project 2018-04635 to JH), the Icelandic Research Fund (Project 206529-051 and #090207021 to KPM).

## Author Contribution Statement

PRM, JH, and KPM conceptualized the study. PRM prepared bioinformatic pipelines. AR, TES, and WW conducted analyses. TES and AR wrote the manuscript with input from all authors. All authors have contributed to the final version of the manuscript.

## Data Archiving

Upon acceptance of the manuscript, raw sequencing data will be uploaded on SRA. Output files will be uploaded on dryad and analysis scripts will be made available by TES on GitHub (https://github.com/RockPtarmigan).

## Research Ethics Statement

The Icelandic rock ptarmigan used in the present investigations were collected specifically for a long-term study on the relationship between population changes and the ptarmigan’s health that has been authorized by the Natural Science Institute of Iceland under law 64/1994, chapter 4, article 7 (http://www.althingi.is/lagas/140a/1994064.html)

## Supplementary Figures

**Supplemental Figure 1.**
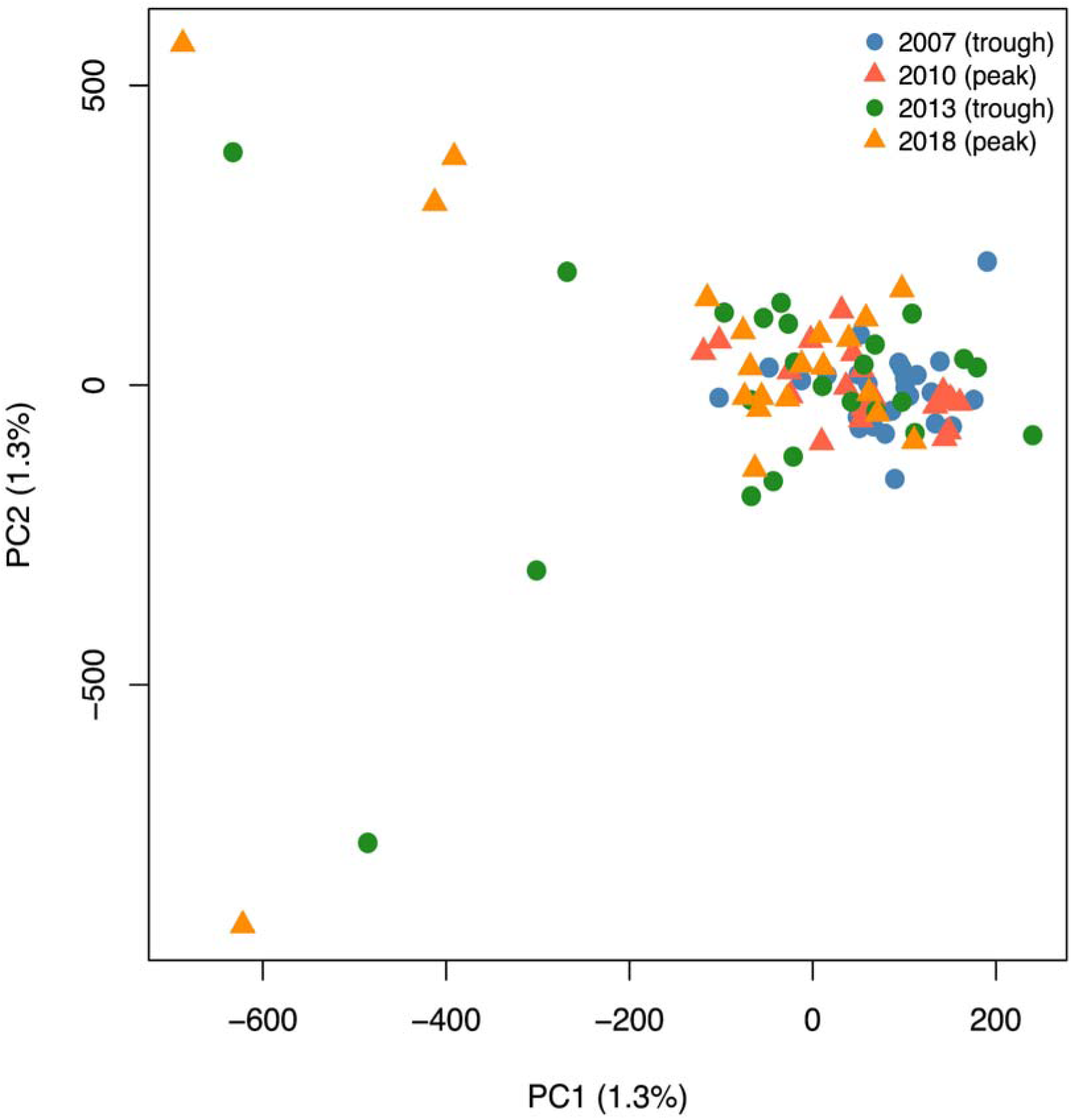
Whole-genome PCA excluding chromosome 16. Principal-component analysis of individual genotypes at the 7,174,144 autosomal SNPs. Chromosome 16 has been excluded as its large inversion polymorphism otherwise dominates the leading PCs. Each point is one of the 91 individuals, colored and shaped by sampling year. Samples do not separate individuals by year, indicating no detectable temporal population structure.

**Supplemental Figure 2.**
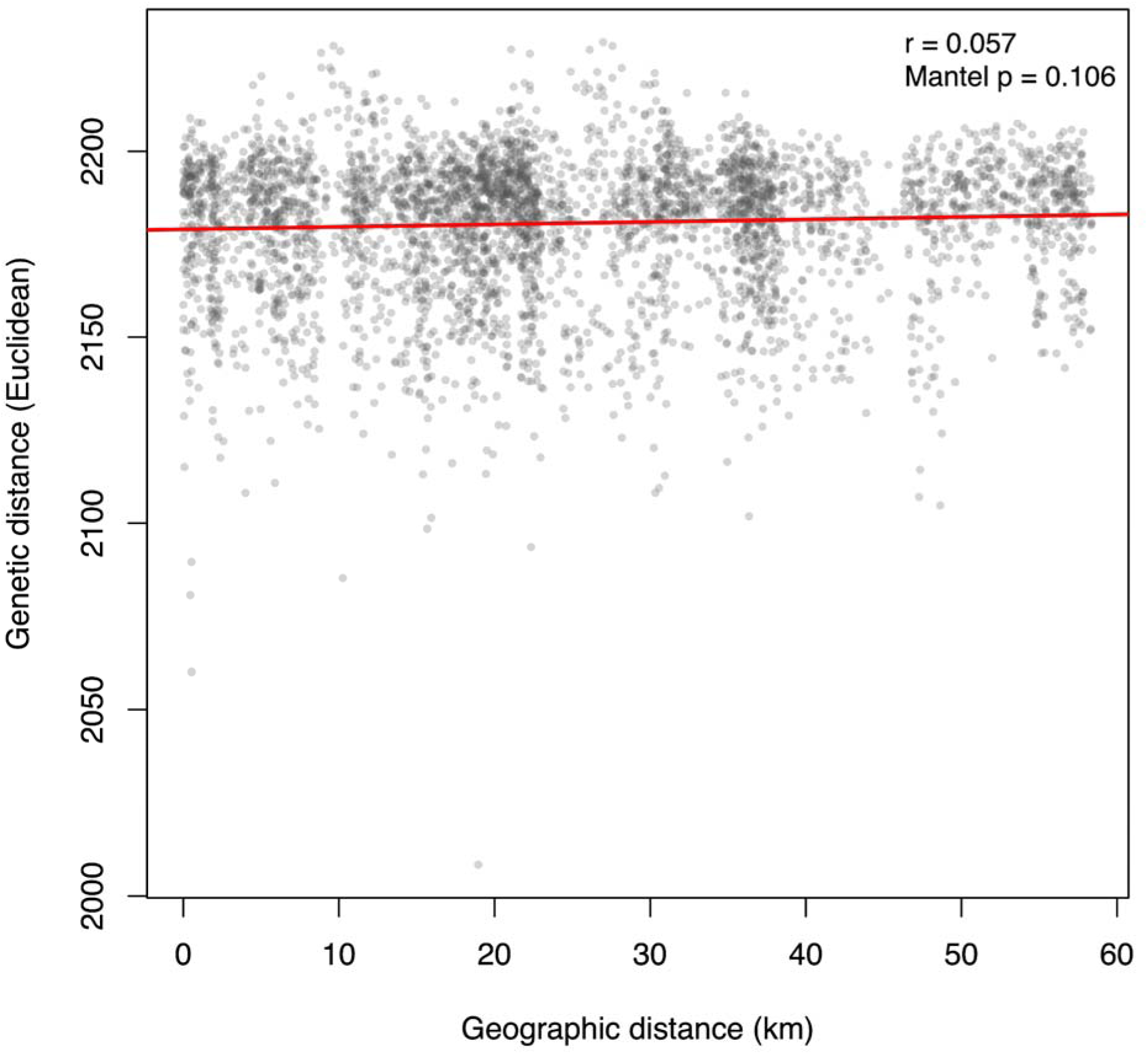
Pairwise genetic distance against geographic distance for all pairs of sampled individuals. Birds were collected within a ∼60 km area of northeast Iceland (pairwise geographic distances 0–58.4 km, median 21.9 km). A Mantel test (Pearson, 9,999 permutations, one-sided) found no significant association between genetic and geographic distance, consistent with a single panmictic population over the sampling area and supporting the pooling of individuals across sites.

**Supplemental Figure 3.**
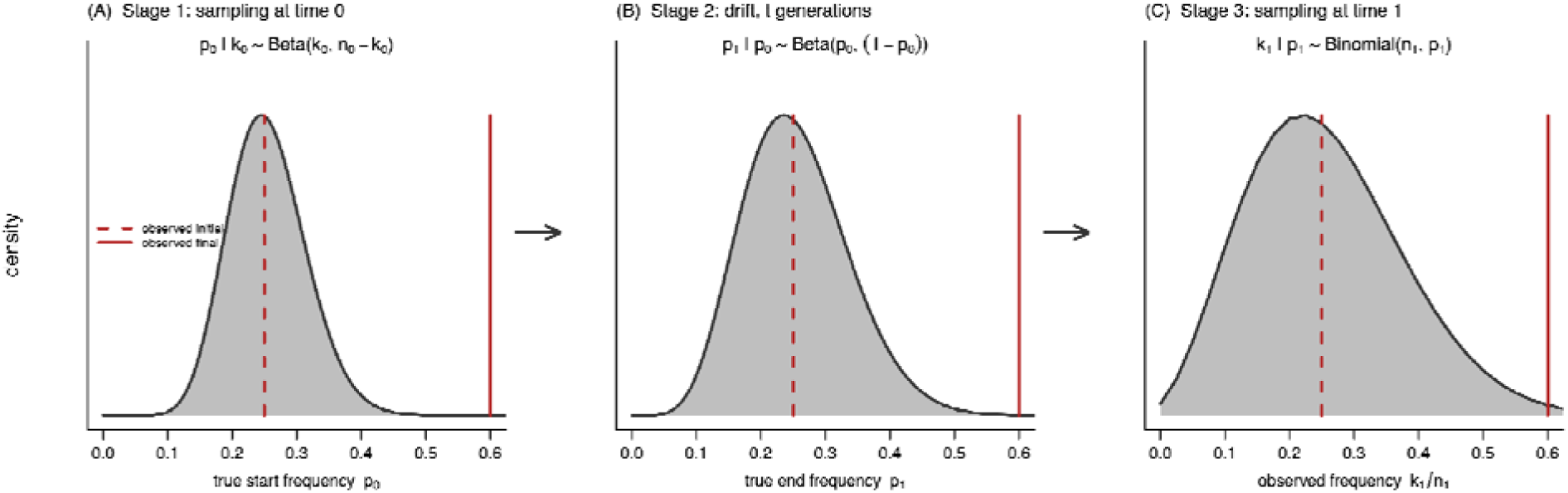
Schematic of the drift-and-sampling null. For each SNP, the observed change in allele frequency between two timepoints is tested against a null built from three compounded sources of variation (left to right). (A) Stage 1: the true initial frequency is uncertain given the finite observed count, represented as a distribution centered near the observed value. (B) Stage 2: genetic drift over t generations broadens this distribution without shifting it on average. (C) Stage 3: sampling of n₁ gene copies at the second timepoint yields the distribution of the observed final frequency, against which the observed change is evaluated. In every panel the dashed red line marks the observed initial allele frequency and the solid red line an example observed final frequency; the p-value is the probability under this null of a change at least as extreme as the one observed. The parameters and distributional forms are illustrative.

**Supplemental Figure 4:**
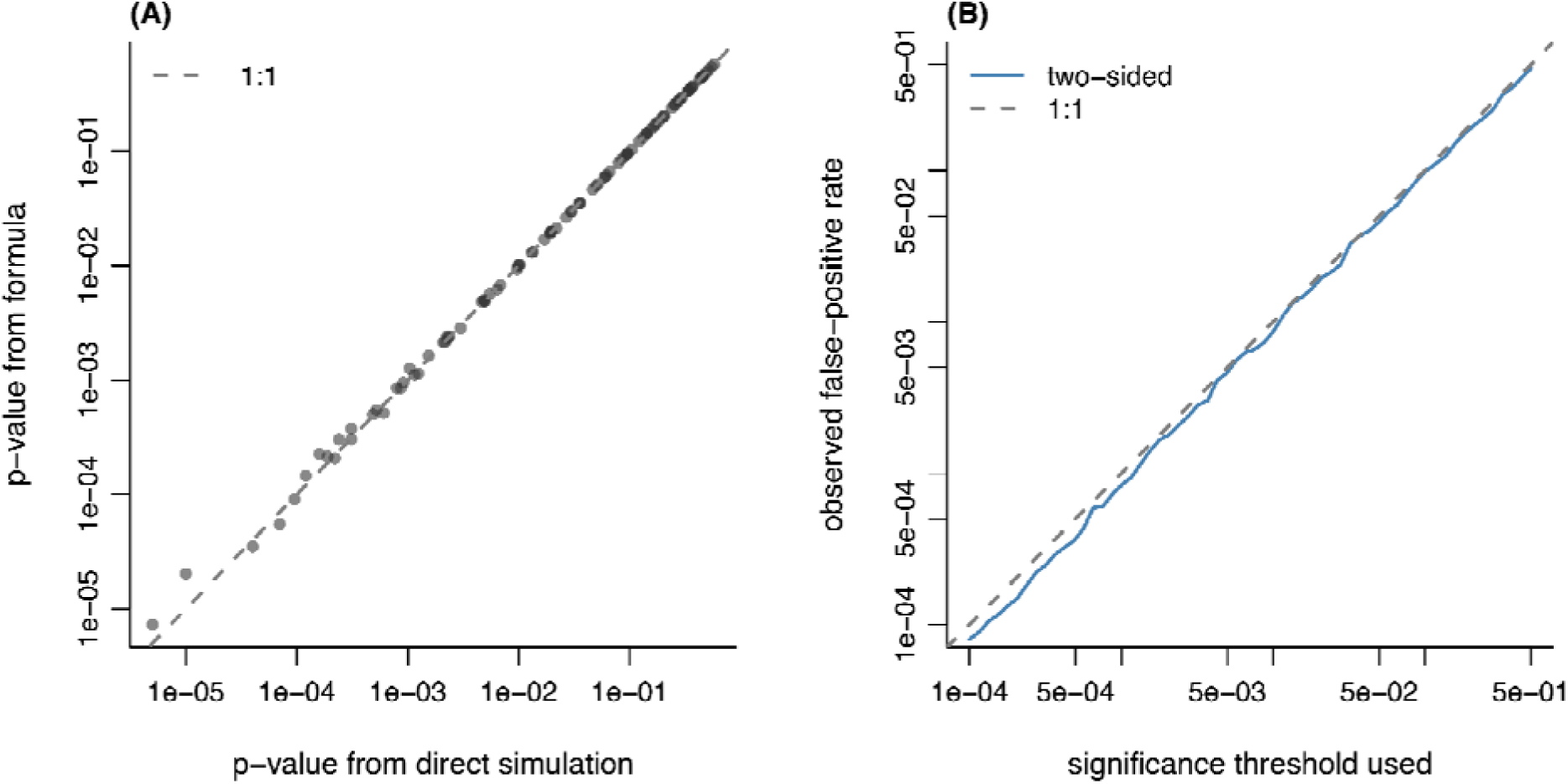
Validation of the closed-form drift-and-sampling test. Both panels use the 2007 to 2010 interval (n_0_ = 52, n_1_ = 40; t = 2 generations; N_e_ = 2,878). For each SNP the observed allele-count change is tested under a three-stage null: uncertainty in the true initial frequency (Jeffreys posterior, p₀ ∼ Beta(k₀+½, n₀−k₀+½)), genetic drift over t generations (beta approximation, p₁|p₀ ∼ Beta(p₀(1−F)/F, (1−p₀)(1−F)/F), with F = 1−(1−1/2NN)N), and binomial resampling at the end (k₁ ∼ Binomial(n₁, p₁)). This compound is evaluated analytically as a moment-matched beta-binomial. (A) Across a grid of observed counts, the closed-form p-value versus the p-value from a direct 200,000-draw Monte Carlo simulation of the same three-stage model; points fall on the 1:1 line (median relative difference 0.3% for p ≥ 0.01), with scatter at the smallest values reflecting the finite resolution of the simulation. (B) False-positive calibration: loci were simulated under the null (no selection) across a range of true starting frequencies and tested with the two-sided (minimum-likelihood) closed-form test; the observed false-positive rate tracks the nominal significance threshold (1:1), confirming the test is well calibrated (marginally conservative at the most stringent thresholds).

**Supplemental Figure 5.**
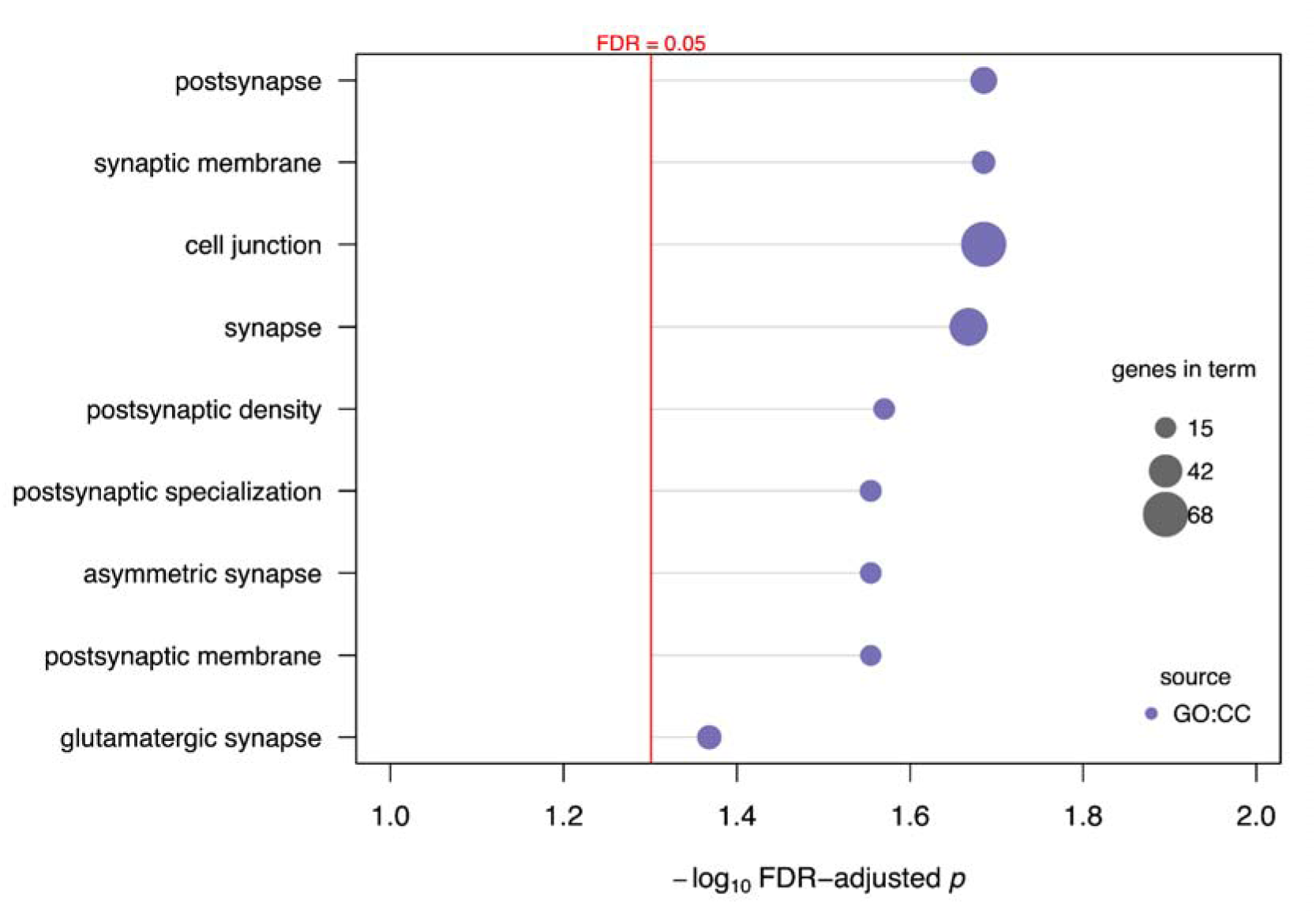
GO enrichment of the top-1% ZFst genes. Genes overlapping the top 1% of pooled peak-versus-trough ZFst windows (537 genes; 411 named protein-coding, tested as Gallus gallus orthologs) were tested for Gene Ontology and pathway enrichment with g:Profiler against all named protein-coding genes genome-wide. Points show the terms passing FDR < 0.05, ordered by significance. Point color indicates the annotation source and point size the number of query genes in the term.

**Supplemental Figure 6.**
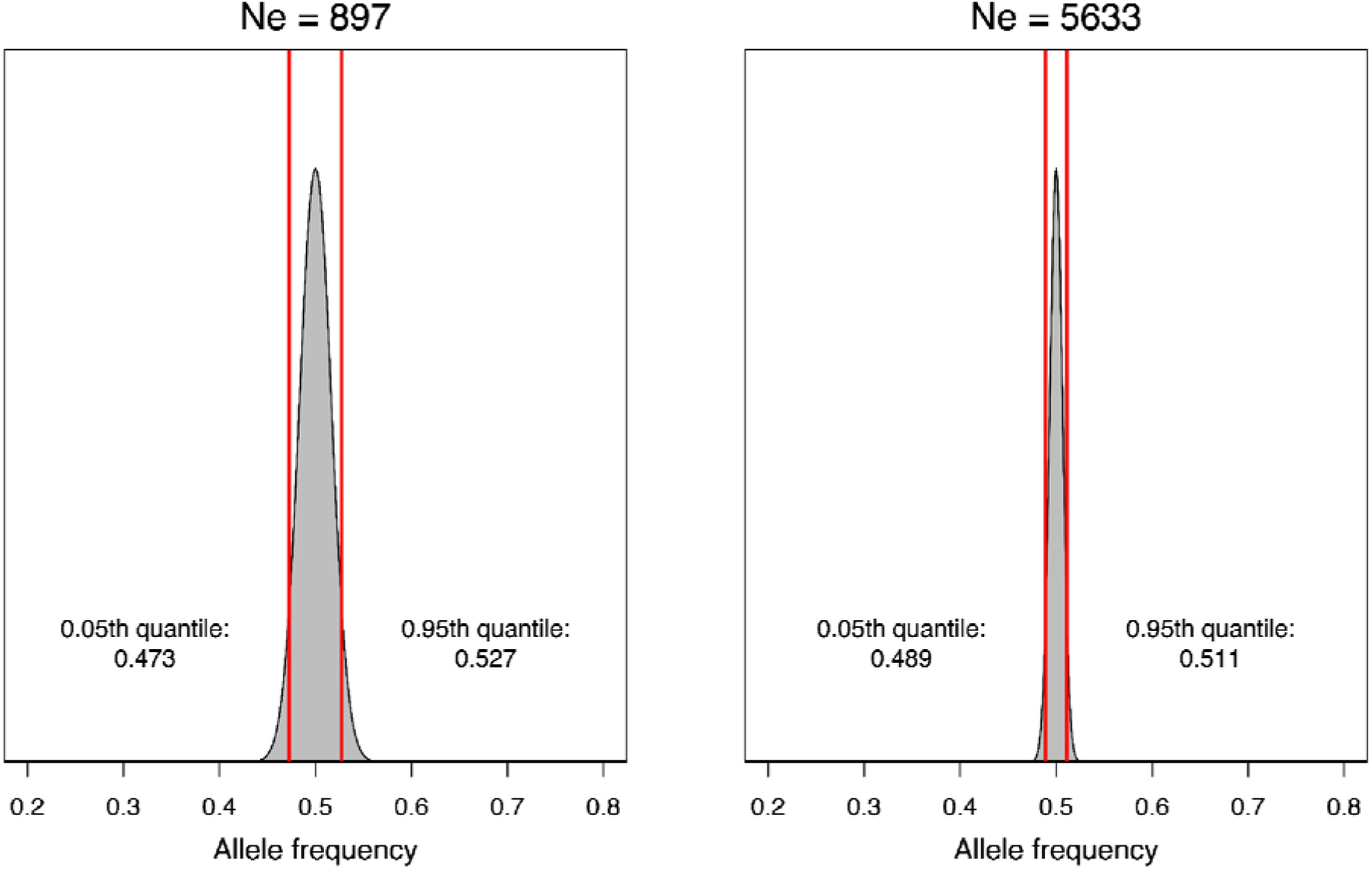
The effects of genetic drift in our populations. Each panel shows the neutral drift expectation for a locus starting at p₀ = 0.5 after t = 2 generations for the range of Ne estimated for all sequential sampled time intervals. Red vertical lines mark the 5th and 95th percentiles. At Ne = 897, the lowest per-interval estimate, drift is expected to shift the allele frequency by at most ≈0.027 in either direction, while at Ne = 5633, the expected shift is smaller still, ≈0.011. Across the full range of estimated Ne, therefore, neutral drift over a single sampling interval moves allele frequencies by only a few percent.

**Supplemental Figure 7.**
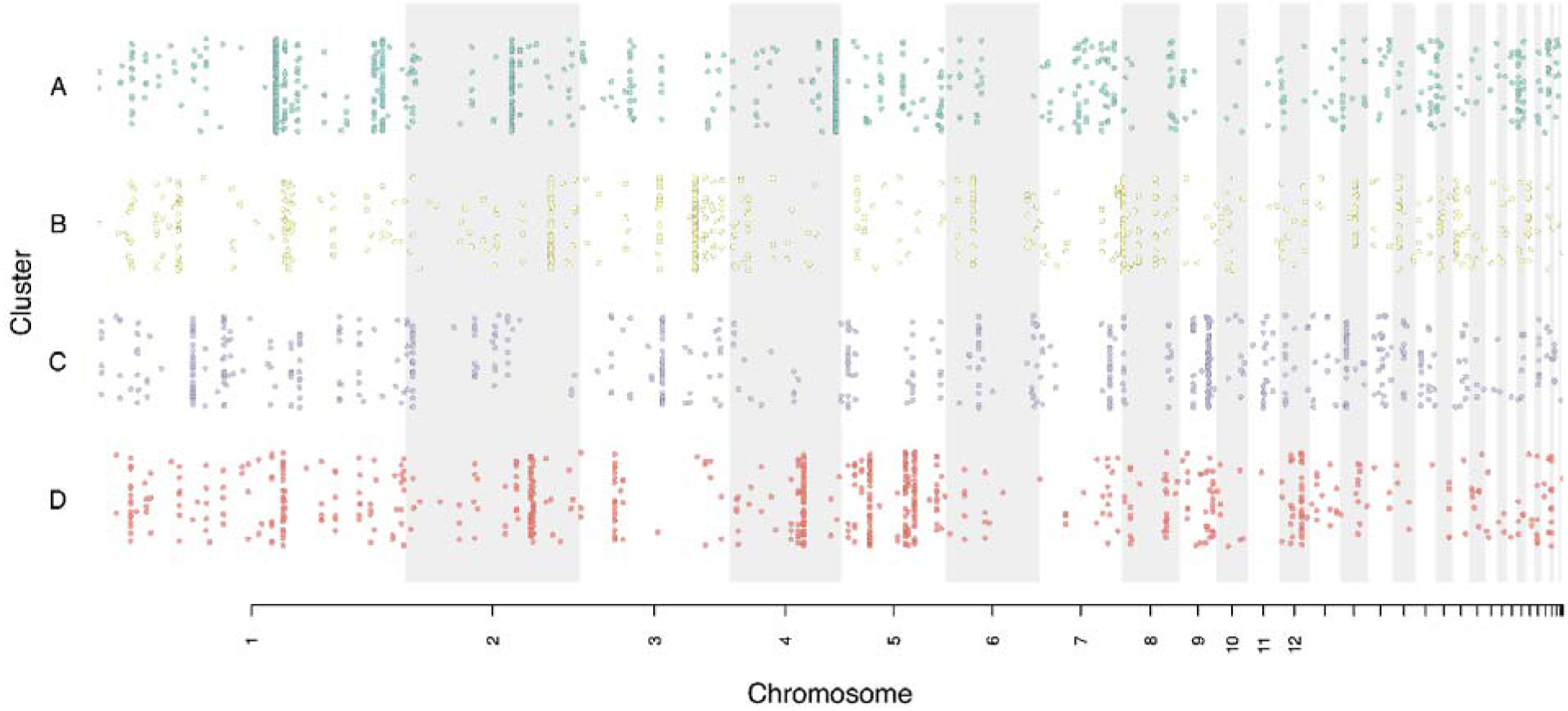
Genomic distribution of the trajectory-cluster SNPs. Each row shows the SNPs assigned to one trajectory cluster (A–D; p < 0.0001 candidate set, clustered at |r| = 0.80), plotted by genomic position and colored by cluster, with points jittered vertically within each row; alternating grey bands mark chromosomes (largest to smallest, labelled beneath). Every cluster is distributed across ≥27 chromosomes rather than being confined to a single region, indicating that the correlated allele-frequency trajectories reflect genome-wide co-fluctuation rather than physical linkage within one haplotype block.

**Supplemental Figure 8.**
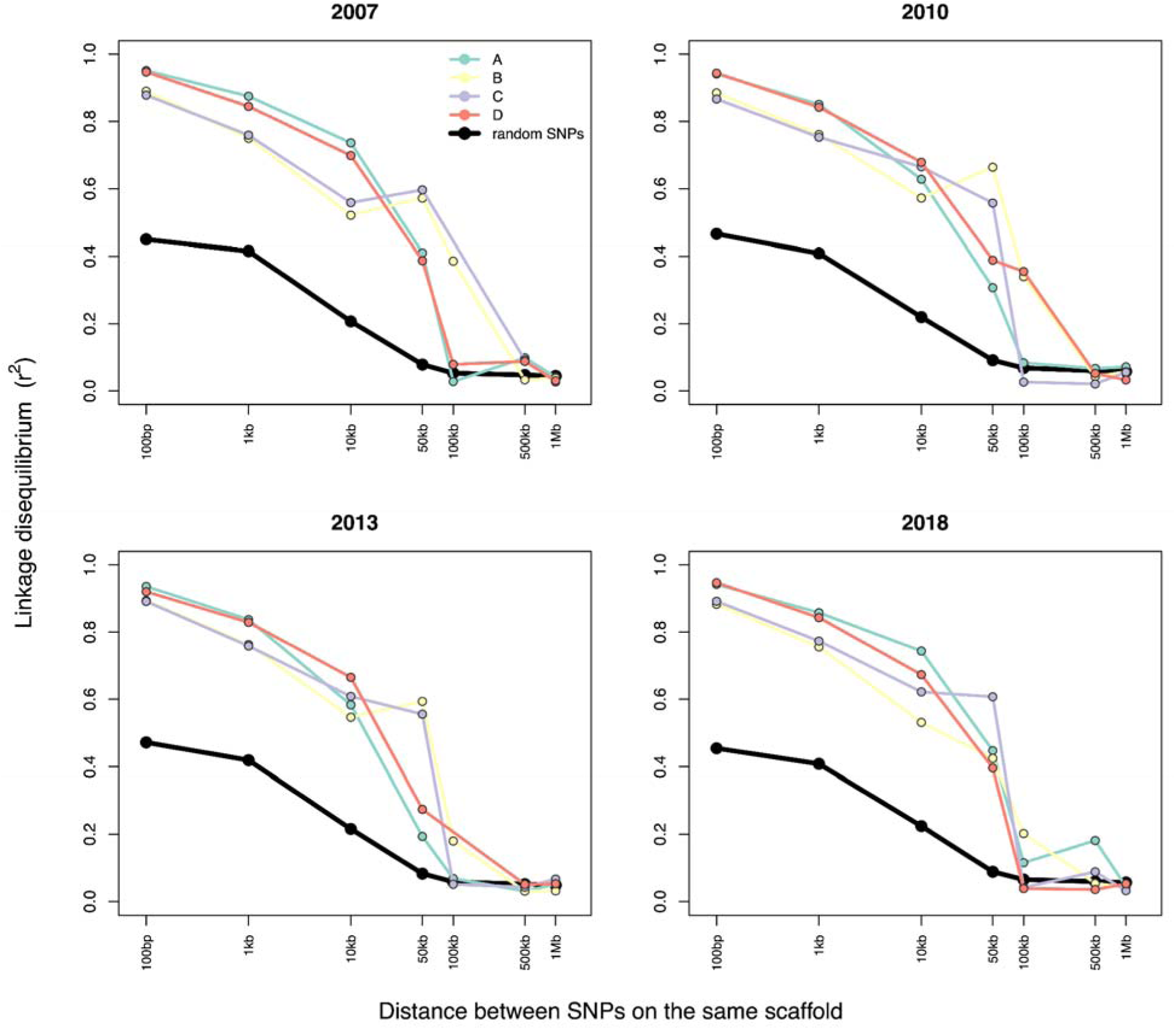
Linkage-disequilibrium decay within trajectory clusters. For each sampling year, genotypic linkage disequilibrium (r²; squared correlation of genotype across individuals, missing calls imputed to the per-SNP mean) is plotted against physical distance between SNP pairs on the same chromosome, averaged within distance bins (bins with fewer than 20 pairs omitted). Colored lines show the four trajectory clusters (A–D; p < 0.0001 candidates, clustered at |r| = 0.80); the black line shows a genome-wide background of 10,000 randomly sampled SNPs. Within every cluster, r² decays to the random-SNP baseline (∼0.05) by ∼100 kb–1 Mb, so cluster members more than 100kb apart are effectively unlinked, confirming that the shared allele-frequency trajectories are not a product of physical linkage.

**Supplemental Figure 9.**
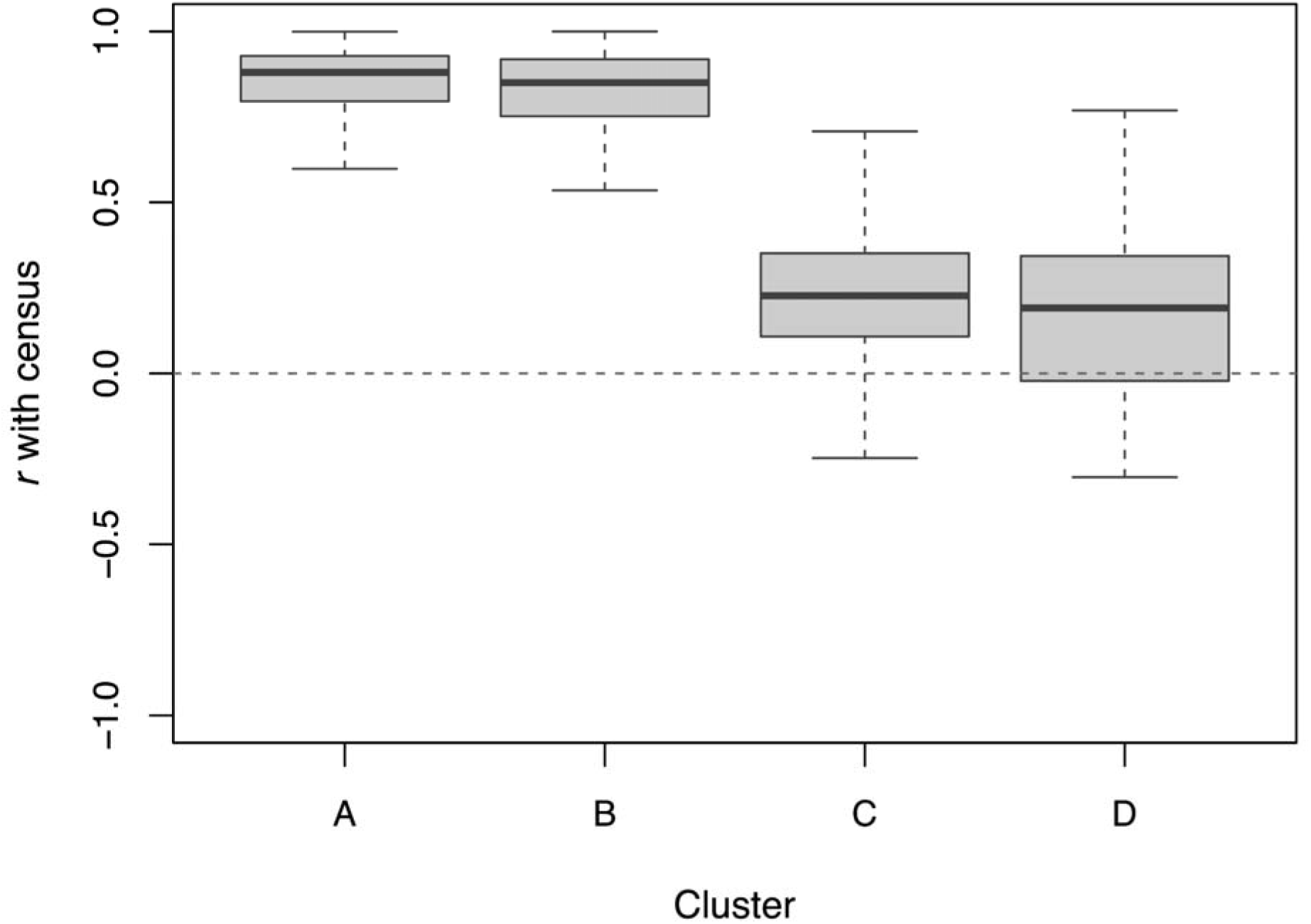
Per-SNP correlation with census size, by trajectory cluster. For each SNP in the trajectory clusters (A–D; p < 0.0001 candidates, clustered at |r| = 0.80), the signed Pearson correlation between its allele-frequency trajectory and the regional census counts across the four sampling years. Each SNP was oriented to align with its cluster mean and each cluster oriented so its mean runs positively with census, so that positive values indicate tracking of census size; the dashed line marks r = 0. Clusters A and B comprise SNPs tightly and positively correlated with census, whereas clusters C and D have little census association. Boxes show the median and interquartile range, whiskers extend to 1.5× IQR, and outliers are omitted.

